# Solution structure of the phosphatidylinositol 3-phosphate binding domain from the *Legionella* effector SetA

**DOI:** 10.1101/2022.02.13.480292

**Authors:** Wendy H.J. Beck, Thais A. Enoki, Xiaochun Wu, Qing Zhang, Linda K. Nicholson, Robert E. Oswald, Yuxin Mao

**Affiliations:** Weill Institute for Cell and Molecular Biology, Cornell University, Ithaca, NY 14853, USA; Department of Molecular Biology and Genetics, Cornell University, Ithaca, NY 14853, USA; Department of Molecular Medicine, Cornell University College of Veterinary Medicine, Ithaca, NY 14853, USA

## Abstract

*Legionella pneumophila* is a facultative intracellular pathogen that causes Legionnaires’ disease or Pontiac fever in humans upon accidental inhalation of *Legionella*-contaminated aerosols. During infection, *L. pneumophila* secretes more than 300 effectors into the host for the biogenesis of a replication-permissive niche, known as the *Legionella containing* vacuole (LCV). Among these, a large number of effectors harbor protein domains that recognize specific phosphoinositide (PI) lipids and mediate the anchoring of these effectors to the surface of LCV or other host membrane-bound organelles. The *Legionella* effector SetA contains a unique C-terminal domain (SetA-CTD) that has been shown to specifically bind with phosphatidylinositol-3-phosphate (PI(3)P) and target SetA to endosomes and LCVs. Here, we report the NMR solution structure of SetA-CTD, which mainly comprises a four α-helix bundle. The structure reveals a basic pocket at one end of the α-helix bundle for PI(3)P binding and two hydrophobic loops for membrane insertion. Mutations of key residues involved in lipid binding result in the loss of SetA in membrane association and endosomal localization. Structural comparison with other PI(3)P-binding domains highlights a general theme applied by multiple families of phosphoinositide-binding domains across species.

## Introduction

Phosphoinositide (PI) lipids are a collection of seven derivatives of phosphatidylinositol reversibly phosphorylated at the 3’, 4’, and 5’ positions on the inositol ring. Although PIs comprise approximately 10% of total lipids within the cell, these lipids play crucial roles in a variety of cellular processes, including cellular signaling, cytoskeleton organization, and membrane trafficking (1–4). Furthermore, PIs also function as a “zip code” for intracellular organelles and define the identity of endomembrane compartments. For example, phosphatidylinositol-4-phosphate (PI(4)P) is enriched primarily in the Golgi apparatus, whereas phosphatidylinositol-3-phosphate (PI(3)P) is primarily enriched in early and late endosomes. The unique distribution of PIs facilitates spatial organization for host proteins that contain one or more PI-binding domains to specific intracellular compartments. Since the first characterization of the pleckstrin homology (PH) domain as a specific PI-binding module (5), there are at least 10 families of PI-binding domains have been documented (6–8). These PI-binding domains adopt a variety of structural folds and apply diverse strategies for PI recognition and membrane association (9,10). PI-binding domains are not restricted to proteins from eukaryotes; new PI-binding modules have frequently been discovered in bacterial pathogens (11–14), particularly in the opportunistic pathogen *Legionella pneumophila* (15–18).

*Legionella pneumophila* is an intracellular bacterial pathogen that is the causative agent of Legionnaire’s Disease, a severe form of pneumonia (19,20). Upon uptake by host cells, *L. pneumophila* delivers over 300 effector proteins into the host cytosol (21,22) for the remodeling of the *Legionella*-containing vacuole (LCV) to enable intracellular proliferation. One key feature of the LCV remodeling process is the dynamic transition of the PI composition on the LCV. Within 1 minute after entry into host cells, PI(3)P accumulates on the LCV and is then gradually replaced by PI(4)P within 2h post-infection (23). This programmed conversion of PIs on the LCV is mediated by a number of PI kinases (24–27) and phosphatases (28–30), as well as phospholipases (31–33), from both the bacterium and the host. The spatiotemporal control of PI composition on the LCV provides a dynamic anchoring platform for a large number of *Legionella* effectors harboring unique domains that specifically bind to PI(4)P and/or PI(3)P (18,34).

Interestingly, most of the PI-binding domains found in *Legionella* effectors have no sequence or structural homology to eukaryotic PI-binding domains. For example, the *Legionella* effector SidM (DrrA) contains a C-terminal P4M domain that has high affinity and selectivity for PI(4)P and mediates the targeting of SidM to the LCV (27). Structural studies revealed that the P4M domain is composed of six helices with three long helices forming a pillar structure (35,36). The top of the pillar, together with other short helices and loops, forms a deep pocket, accommodating the head group of a PI(4)P molecule. Hydrophobic residues aligned along the rim of the pocket function as a “membrane insertion motif” (MIM), contributing to the high affinity for P4M membrane association (36). Another example is the P4C domain from the *Legionella* effector SidC and its paralog SdcA, which specifically recognizes PI(4)P and facilitates the LCV localization of SidC (37,38). Structural and biochemical characterization revealed that the P4C domain is comprised of a four α-helical bundle and one end of the bundle, together with two loops enriched with hydrophobic residues, forms a PI(4)P-binding pocket (39). Besides the well-characterized P4M and P4C domains, a long list of *Legionella* effectors has been shown to possess unique PI-binding modules. A recent global bioinformatics study identified three families of novel PI(3)P-binding domains presented in 14 *Legionella* effectors (17). Despite the rapid expansion of the families of PI-binding domains, our understanding of the molecular mechanism involved in phosphoinositide recognition and selective membrane targeting is still limited due to the lack of structural information of these prokaryotic PI-interacting domains.

In this study, we characterized a specific PI(3)P-binding domain from the *L. pneumophila* effector, SetA (lpg1978). SetA possesses an N-terminal mono-*O-*glucosyltransferase domain and a C-terminal domain responsible for PI(3)P-binding with high affinity and specificity (40,41). Recent studies showed that SetA mediates the posttranslational glucosylation of a large array of host factors, including actin, vimentin, the chaperonin CCT5, and the small GTPase Rab1a (42–44). Our recent data showed that SetA modifies the transcription factor TFEB and causes TFEB nuclear translocation and activation (45). SetA localizes to the LCV and endosomal membrane via its C-terminal PI(3)P-binding domain (40); however, the mechanism of PI(3)P binding has yet to be resolved. Here, we present the NMR solution structure of the PI(3)P-binding domain of SetA. The structure reveals that the PI(3)P-binding domain of SetA comprises four α helices and a two-stranded β sheet. The four α helices form a helical bundle, while the two β strands are inserted within the loop connecting helix α1 and α2. Our structure further reveals a potential PI(3)P-binding pocket at end of the α helical buddle and the structural determinants responsible for PI(3)P binding were characterized by biochemical and cellular assays. Our findings provide structural insights into intracellular targeting of SetA and further shed light on the molecular mechanism that governs the recognition of PIs by novel PI-interacting domains from prokaryotic species in general.

## Results

### NMR Structural determination of the SetA PI(3)P-binding domain

SetA was previously characterized as a glucosyltransferase. It contains an N-terminal catalytic domain, which possesses a canonical DxD catalytic motif and transfers a glucose moiety to its targets from UDP-glucose, and a C-terminal domain (CTD: residues 507-644) that is capable of PI(3)P-binding and is essential for the targeting of SetA to endosomes of host cells (40) (Figure 1A). Sequence homology analysis revealed that the SetA-CTD has a unique sequence conserved only on SetA-related proteins (Supplemental Figure 1). Furthermore, the structural homology search yielded no significant hits. To gain insight into the molecular mechanism of PI(3)P binding and endosomal targeting, we carried out structural studies of SetA-CTD. Initial crystallization attempts of this domain were unsuccessful; therefore, we used 3D NMR spectroscopy for structural determination. We initially tested with a SetA construct spanning residues 507-644; however, the ^1^H-^15^N HSQC spectrum corresponding to this construct was dominated by strong peaks characteristic of unstructured peptides. We subsequently generated a series of truncations of the SetA CTD for NMR spectroscopic analysis. Of the constructs tested, the SetA truncation encompassing residues 523-629 yielded well-dispersed peaks in the ^1^H-^15^N HSQC spectrum (Supplemental Figure 2). This fragment of CTD retains the PI(3)P binding capability as shown below and thus was used for further NMR spectra acquisition and structural determination.

**Figure 1.**
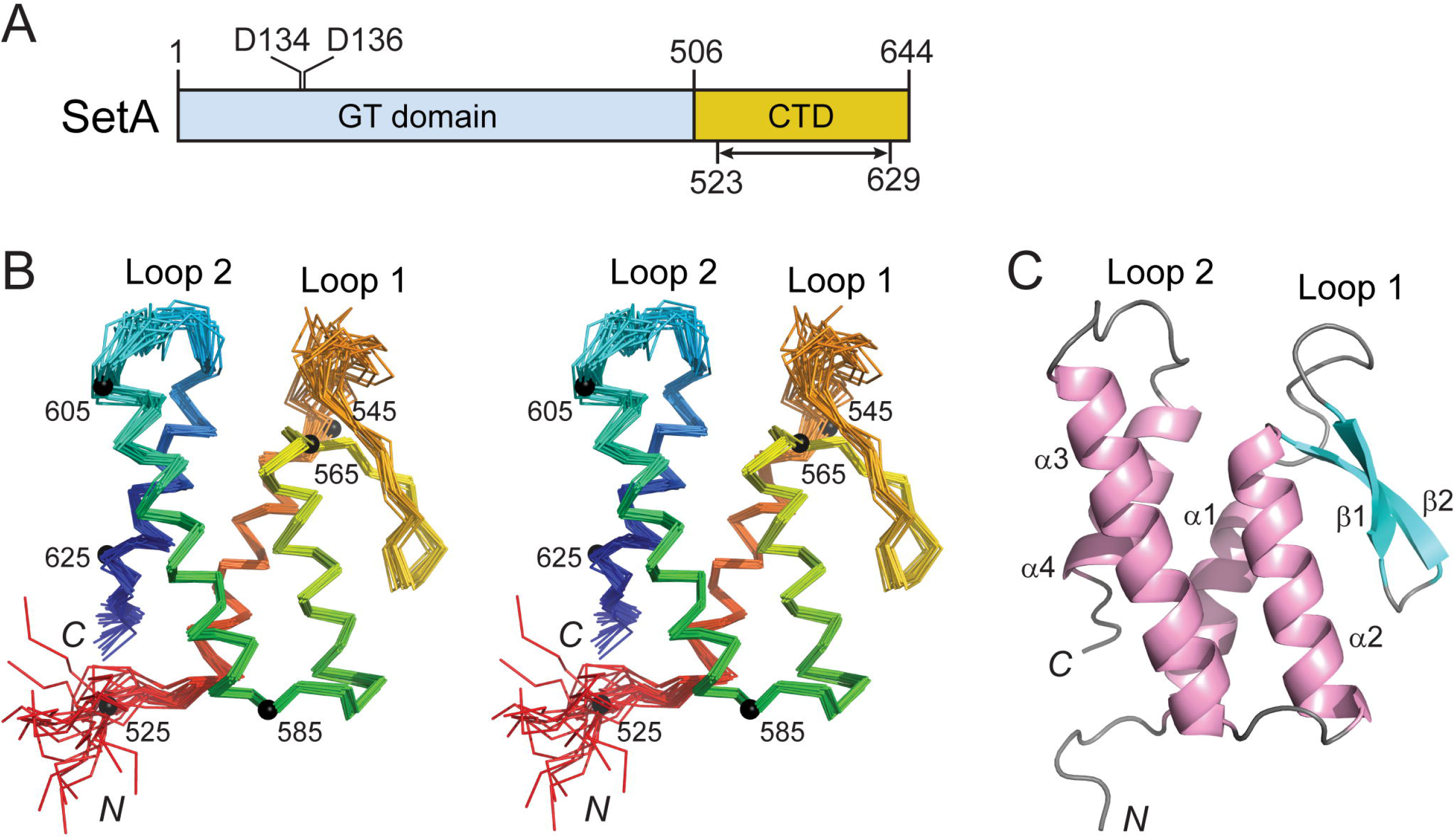
NMR structure of the C-terminal domain of SetA. (A) Schematic diagram of the domain structure of SetA. SetA contains an N-terminal glycosyltransferase (GT) domain (colored in cyan) and C-terminal PI(3)P binding domain (CTD) (yellow) (B) A stereoview of superimposed C^α^ backbone traces of the 20 lowest-energy calculated structures. Traces are rainbow-colored with the N-terminus in red and the C-terminus in blue. Numbers and corresponding small filled circles indicate residues multiple of 20. (B) Ribbon diagram of a representative structure from (A) shown in the same orientation.

As a first step towards structure determination, we collected a series of 3D NMR spectra for chemical shift assignment (see details in methods) and the ^1^H, ^13^C, and ^15^N NMR chemical shifts for residues in this fragment were assigned to near completion. Tertiary structures were calculated using the Ponderosa-C/S server (46,47) based on a total of 2661 NOE-derived distance restraints, 153 dihedral angle constraints, and 106 hydrogen bond restraints. The final ensemble of 20 calculated structures is presented in backbone C^α^ trace as a stereo pair (Figure 1B) and structural statistics are listed in Table 1. In agreement with the distribution of the NOE constraints along with the peptide (Supplemental Figure 3), the structure of regions containing regular secondary structure (α helix and β strand) is highly converged whereas the N-terminus and two long loops (Loop1 and Loop2) connecting the α helices are flexible (Figure 1B). The flexibility of the two loops may correlate with their function in membrane binding (see below).

**Table 1.** NMR and structural refinement Statistics for SetA.

|  |  |
| --- | --- |
| **NMR distance and dihedral constraints** |  |
| NOE distance restraints | 2661 |
| Short range ($\vert i-j\vert \leq 1$) | 1542 |
| Medium range ($1 < \vert i-j\vert \leq 4$) | 562 |
| Long range ($\vert i-j\vert > 4$) | 451 |
| Dihedral angle constraints |  |
| $\phi$ | 71 |
| $\psi$ | 82 |
| Hydrogen bond restraints | 106 |
| Disulfide bond | 0 |
| **Structure statistics** |  |
| Residual NOE violations (mean ± SD) | |
| Number of distance constraints (0.1 - 0.2 Å) | 21.32 ± 4.75 |
| Number of distance constraints (0.2 - 0.5 Å) | 4.18 ± 1.76 |
| Maximum violation (Å) | 0.392 ± 0.045 |
| RMS violation (Å) | 0.021 ± 0.001 |
| Residual dihedral angle violation (mean ± SD) | |
| Number of dihedral angle constraints (1- 5°) | 1.68 ± 0.99 |
| Maximum violation (°) | 1.57 ± 0.48 |
| RMS violation (Å) | 0.19 ± 0.05 |
| Deviation from idealized geometry |  |
| Bond (Å) | 0.011 |
| Angle (°) | 1.5 |
| Average pairwise rmsd (Å) |  |
| Heavy | 0.6 <sup>a</sup> |
| Backbone | 1.0 <sup>a</sup> |
| Ramachandran plot statistics (%) |  |
| Residues in most favored regions | 88.9 |
| Residues in additionally allowed regions | 9.5 |
| Residues in generously allowed regions | 1.7 |
| Residues in disallowed regions | 0 |

<sup>a</sup>Pairwise rmsd was calculated from 20 structures aligned on well defined secondary structure elements: Residues: 8 - 23, 32 - 81, 92 - 106

### The overall structure of the PI(3)P-binding domain of SetA

The NMR structure of SetA PI(3)P-binding domain (a.a. 523-629) mainly consists of an anti-parallel four α-helical bundle with a two-stranded β sheet inserted into the loop (Loop1) connecting helices α1 and α2 (Figure 1C). Interestingly, a pocket is formed between Loop1 and Loop2 at one end of α-helical bundle (Figure 1C and 2). This pocket has an overall positive electrostatic surface potential due to the presence of residue R612, which resides at the bottom of the pocket (Figure 2A and B). In addition, both rims of the pocket are lined with bulky hydrophobic residues: L549 and L550 in Loop1 and F608 and F609 in Loop2 (Figure 2B). These structural observations suggest that this pocket is likely the PI(3)P binding site with the positively charged R612 accommodating the negatively charged 3’ phosphate on PI(3)P and the hydrophobic residues located at the rims of the pocket as the putative MIM.

**Figure 2.**
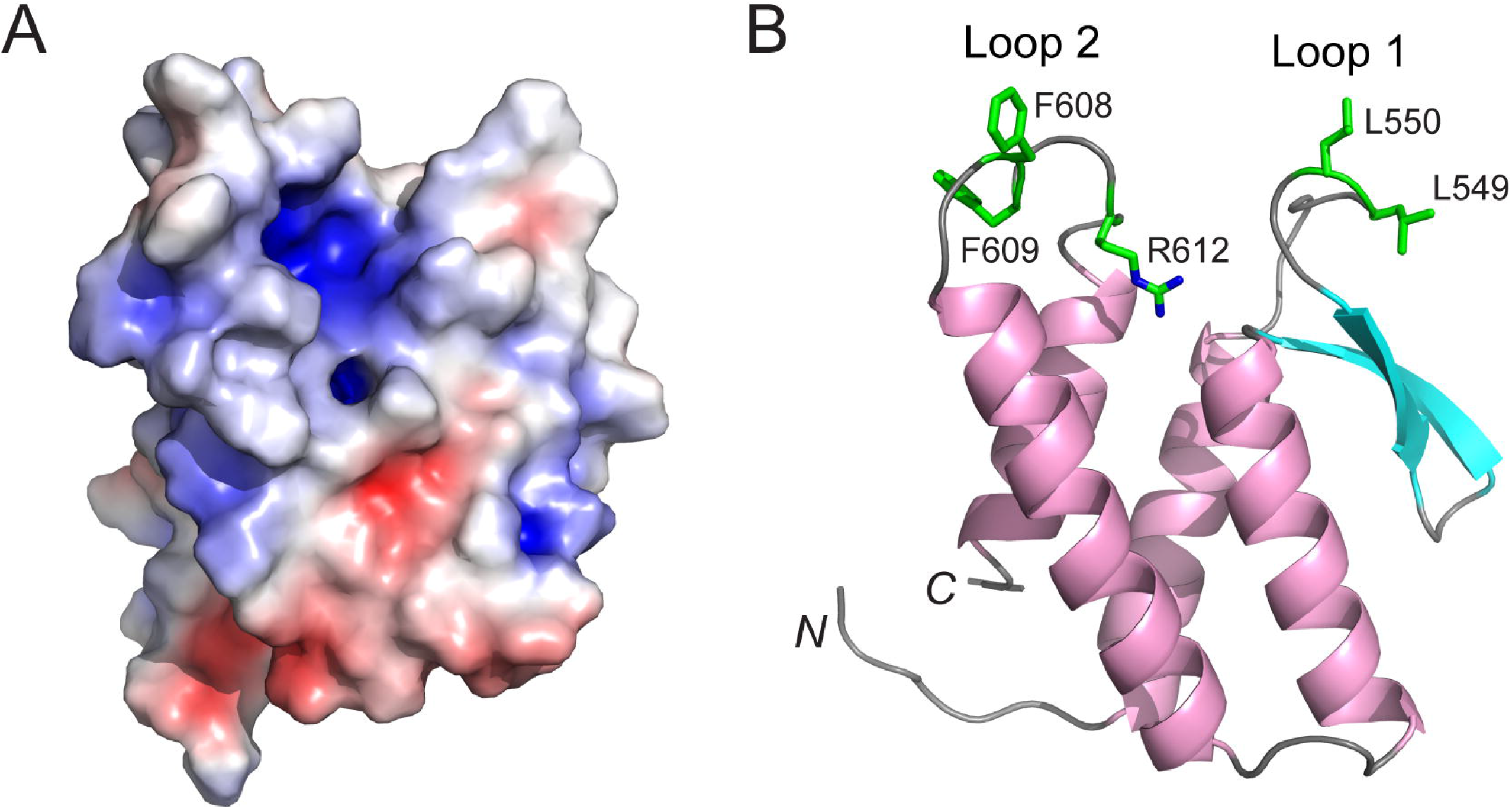
Determinants for PI(3)P binding and membrane association in SetA-CTD. (A) Molecular surface of SetA-CTD. The surface is colored based on electrostatic potential with the positively charged region in blue (+5 kcal/electron) and negatively charged surface in red (−5 kcal/electron). (B) Ribbon diagram of SetA-CTD. Residues that play a role in PI(3)P binding and membrane interaction are highlighted in sticks. R612 resides at the bottom of a pocket is likely to recognize the PI(3)P headgroup. Loop 1 (L549 and L550) and Loop 2 (F608 and F609) residues are shown in sticks and colored in green. These residues form the membrane interacting motif (MIM).

### Membrane interaction by the SetA PI(3)P-binding domain

To characterize the membrane interaction by the SetA PI(3)P-binding domain, we performed fluorescence microscopy imaging-based in vitro assays. Recombinant proteins of SetA-CTD fused with an EGFP were prepared and incubated with giant unilamellar vesicles (GUVs) containing POPC, POPS, with or without PI(3)P. In agreement with previously reported PI specificity for SetA (40), strong GFP signals were observed on the surface of PI(3)P-containing GUVs, whereas in the absence of PI(3)P, no GFP signal was detected (Figure 3). Next, to test whether the bulky hydrophobic residues aligned at the rims of the pocket are important for SetA membrane binding, we mutated these hydrophobic residues to a smaller polar residue, serine. Membrane binding by either the Loop 1 mutant (L549S/L550S), or the Loop 2 mutant (F608S/F609S), or the Loop 1&2 mutant carrying all four substitutions was abolished (Figure 3), suggesting these hydrophobic residues are essential for membrane binding by the SetA PI(3)P-binding domain and likely function as the MIM. We further examined the role of the positively charged residue R612 in PI(3)P binding. As expected, the SetA R612Q mutant lost the binding to PI(3)P containing GUVs (Figure 3), suggesting that R612 is likely involved in PI(3)P binding by electrostatic interactions with the negatively charged phosphate group on PI(3)P.

**Figure 3.**
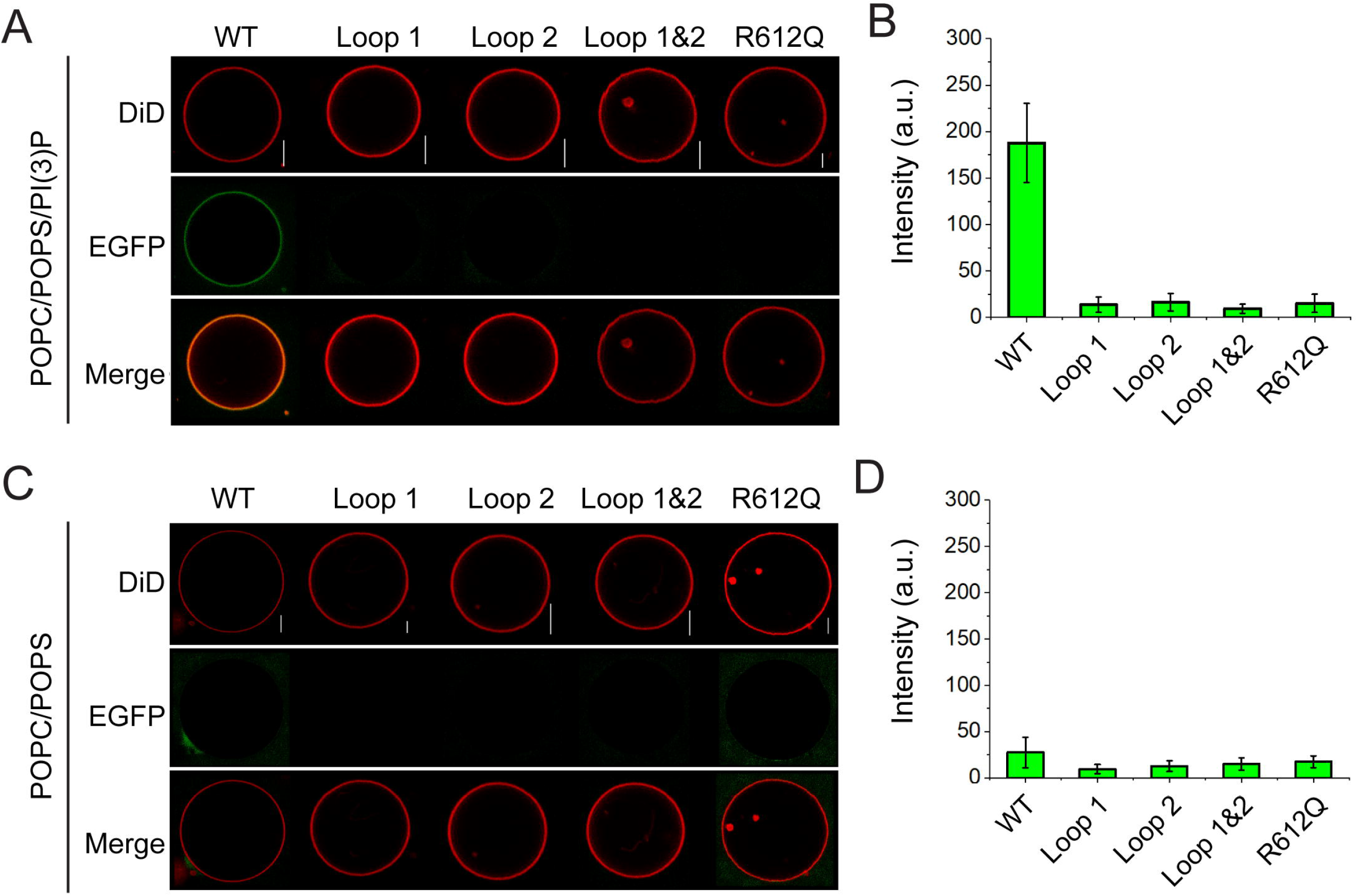
Membrane interaction by the PI(3)P-binding SetA-CTD. (A) Purified recombinant proteins of GFP-SetA-CTD and indicated mutants were incubated with giant unilamellar vesicles (GUV) formed with POPC/POPS/PI(3)P. The proteins bound to the GUV were monitored by fluorescence microscopy. (B) Quantification of the EGFP signal intensity on the GUVs. (the intensity is averaged from 3 independent measurements?) (C) GUV binding assay of wild type and mutant GFP-SetA-CTD proteins with GUV formed by POPC/POPS. (D) Quantification of the EGFP signal intensity on the GUVs. The error calculated from the binding assay represents the standard error from the measurement of 10 GUVs. Scale bar: 5μm.

SetA has previously been shown to localize on the surface of LCV and endosomes when exogenously expressed (40,41,45). We next determined whether the SetA mutants, which abrogated PI(3)P binding in our in vitro GUV binding assays, would also be defective in targeting SetA to the PI(3)P-positive compartments in the cell. mCherry-tagged SetA-CTD constructs expressing either the wild type PI(3)P-binding domain or the indicated mutants were co-expressed with the PI(3)P probe, EGFP-2xFYVE in HeLa cells. The wild type SetA-CTD displayed a high degree of co-localization with EGFP-2xFYVE (Figure 4A) with about 90% of the SetA signal colocalized with the FYVE-positive puncta (Figure 4B). However, each of the loop mutants carrying the same amino acid substitutions as described above showed a diffused localization. These findings indicate that the hydrophobic residues are essential for the anchoring of SetA to PI(3)P-containing membrane compartments, consistent with the results from the in vitro GUV binding assay (Figure 3). Similarly, the R612Q mutant also displayed an increased cytosolic location and a significant reduction of co-localization with EGFP-2xFYVE (Figure 4). Together, our data demonstrate that the anchoring of SetA to PI(3)P-containing membrane compartments require both a cationic residue, that contributes to the recognition of the headgroup of PI(3)P, and the hydrophobic MIM, which reinforces the association of the protein with the membrane bilayer.

**Figure 4.**
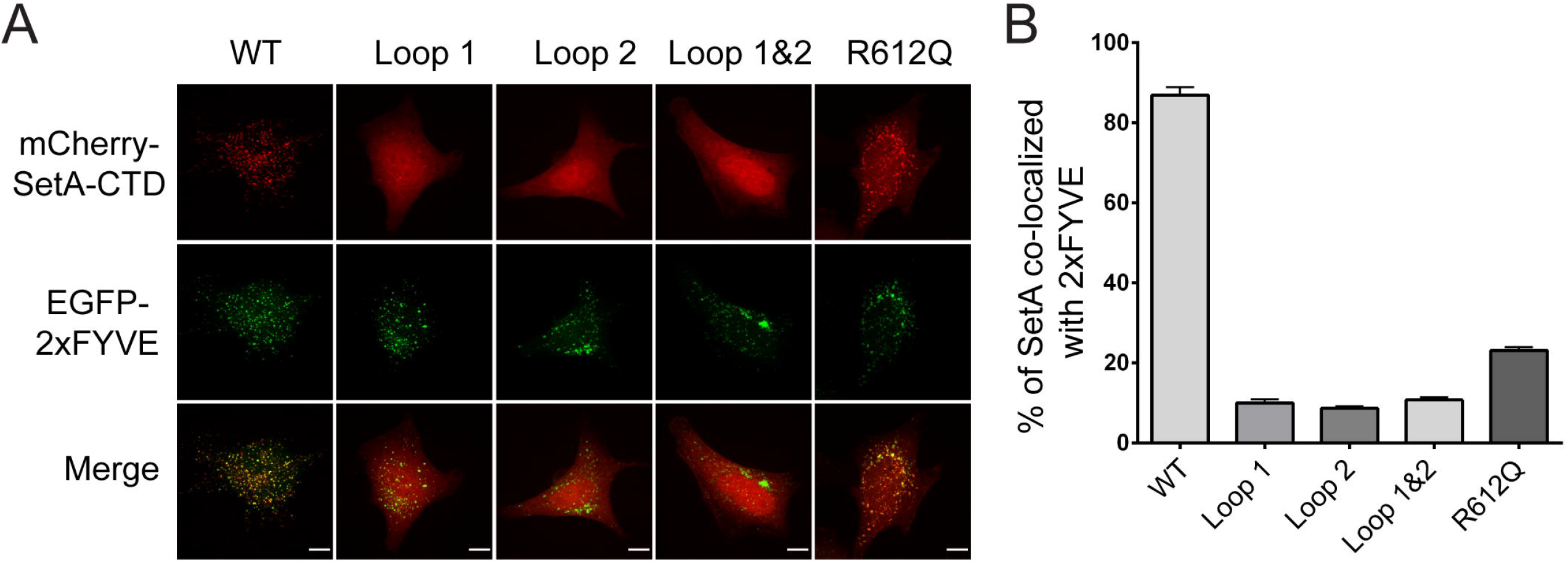
Intracellular localization of mCherry-tagged SetA-CTD. (A) Confocal fluorescent images of HeLa cells cotransfected with EGFP-2xFYVE and mCherry-tagged wild type or indicated SetA-CTD mutants. Scale bar: 10 μm. (B) Quantification of the co-localization percentage of mCherry-SetA-CTD with GFP-2xFYVE. The percentage of colocalization was calculated using the ImageJ software. Values are the mean ± SEM. (n = 10).

### Comparison with other Pl(3)P-binding domains

A wide variety of PI-binding domains have been documented (1). Among these, the FYVE domain (48,49) and a subset of the PX domain (50,51) have been characterized as PI(3)P-specific binding modules. A comparison of SetA-CTD with these eukaryotic PI(3)P binding domains reveals vast differences in the structural fold and the diverse mode of phosphoinositide binding (Figure 5). The SetA PI(3)P-binding domain adopts a four α-helical bundle structure. A PI(3)P-binding pocket is formed by loops connecting the α-helixes with a positively charged Arg residue aligned at the bottom of the pocket. However, the FYVE domain consists of two double-stranded antiparallel β sheets and a short C-terminal α-helix. These secondary structural elements are tethered together by coordinating with two Zn^2+^ ions (48,49,52). A conserved basic motif (RR/KHHCR) spanning across the first β strand forms the positively charged PI(3)P-binding site. On the other hand, the structural core of the PX domain has an N-terminal twisted three-stranded β sheet and a C-terminal cluster of α-helixes, that pack on one side on the β sheet (50,53). Residues from two long α-helixes, the connecting loop between the two α-helixes, and the tip of the β sheet form a pocket that accommodates the headgroup of PI(3)P. Another difference among these three PI(3)P-binding modules resides at their MIMs. In the C-terminal domain SetA, the MIM consists of four bulky hydrophobic residues located at the tip of the two loops. This feature suggests that hydrophobic interaction mediated by the insertion of the MIM residues into the membrane bilayer makes a major contribution to the affinity of membrane association. By contrast, the FYVE domain has fewer MIM residues (in the case of EEA1, only one valine residue), hence has a lower affinity for membrane association and FYVE domain-containing proteins tend to require additional mechanisms, such as dimerization, for membrane association (Figure 5B) (52,54). In the PX domain, although not conserved, large exposed hydrophobic residues are present at protruding loops near the PIP-binding pocket and contribute to membrane binding by membrane penetration (Figure 5C) (50). In summary, despite their markedly different structural folds, the comparison of these PI(3)P-binding domains further suggests that properly positioned positively charged residues and membrane penetrating hydrophobic residues are consensus requirements for their association with PI(3)P-containing membrane.

**Figure 5.**
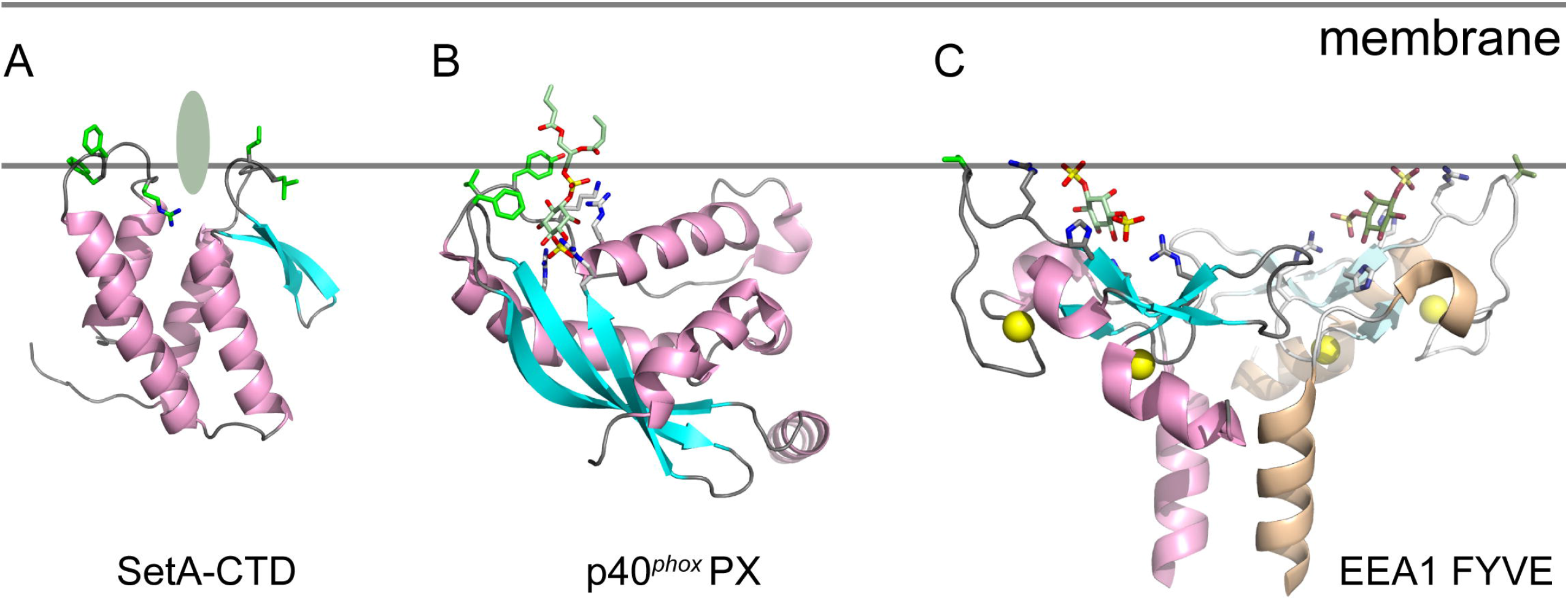
Structural comparison of PI(3)P-binding domains. (A) Ribbon representation of the NMR structure of SetA-CTD. The structure is orientated with the predicted PI(3)P binding site (indicated by a pale green oval) interfaced with the membrane. Residues tested to interact with the membrane are colored in green and shown in sticks. (B) Membrane interaction model of the p40^phox^ PX domain(PDB ID: 1h6h). The bound di-butyl-PI(3)P and membrane interacting residues are shown in sticks. (C) Membrane interaction model of the EEA1 FYVE domain (PDB ID: 1joc). In this model, the FYVE domain is presented as a homodimer with a truncated EEA1 coiled-coil region. The bound inositol-1,3-bisphosphate and membrane interacting residues are shown in sticks.

## Discussion

In this study, we have determined the solution structure of a specific PI(3)P-binding domain from a *Legionella* effector SetA. Our structure reveals a novel PI(3)P-binding domain with a four-alpha-helix-bundle fold. Our structure also predicts that a cationic pocket formed by two flexible loops at one end of the helix bundle serves as the PI(3)P-binding site. Our in vitro liposome binding and intracellular localization assays further validate the structural determinants of SetA-CTD for membrane interaction. Our data highlight a general mechanism of membrane association by PIP-binding modules. These small structural domains are often observed to have a positively charged pocket, which is primarily responsible for the binding with the head group of phosphoinositides and confers the lipid selectivity, and hydrophobic MIM residues, that penetrate the hydrophobic membrane bilayer and contribute largely to the membrane-binding affinity.

Interestingly, PIP-binding modules function beyond anchoring effector proteins to specific membrane-bound organelles, these modules are often observed to regulate the enzymatic activities encoded in other portions of the proteins. For example, *Legionella* effectors SidC/SdcA harbors an N-terminal ubiquitin ligase (SNL) domain and a C-terminal PI(4)P-binding P4C domain. It has been observed that the P4C domain packs against the active site of the SNL domain and inhibits the ubiquitin ligase activity of SidC/SdcA. The binding of the P4C domain to PI(4)P-containing membrane relieves the inhibitory effect and significantly stimulates ubiquitin ligation catalyzed by the SNL domain (39). Similarly, it has been shown that the binding of PI(3)P by the SetA-CTD substantially enhances the glucosyltransferase activity mediated by the N-terminal GT domain of SetA (42). Although the mechanism of activation is not fully understood, our structural results will provide a solid foundation for future investigation of the role of PI(3)P-binding by SetA-CTD.

Results from this study bring forth a potential strategy to develop a novel PI(3)P probe from a prokaryotic origin. The current popular PI(3)P probe is based on the FYVE domain of Hrs (the receptor tyrosine kinase substrate) designed in a tandem fashion (55). However, PIP probes derived from eukaryotic PIP binding modules often show limitations, such as low affinity for targeting lipids or interactions with other eukaryotic proteins. The SetA-CTD appears to have a high specificity and affinity for PI(3)P (40). Its prokaryotic origin may also reduce the interference by other eukaryotic factors. Although more study is warranted to optimize this novel probe, our data suggest that the SetA-CTD has a promising potential to be developed into an accurate and sensitive PI(3)P probe. Furthermore, the SetA-CTD is responsible for the anchoring of SetA to endosomes and the LCV. Deletion of the SetA C-terminal domain causes diffused cytosolic localization of the protein (40,45). These observations suggest that SetA-CTD can also be developed as a fusion tag to target any fused protein of interest to endosomes, where PI(3)P is mostly enriched in the cell.

## Materials and Methods

### Cloning and Site-Directed Mutagenesis

The DNA fragment of SetA-CTD (residue 523-629) was PCR amplified from a plasmid containing the full-length SetA cDNA insert (45). For recombinant protein expression, SetA-CTD was cloned into a pET28(a) fused with a His-Sumo using the BamHI and XhoI cut sites. For mammalian cell expression, SetA-CTD was ligated into the pEGFP-C1 and the pmCherry-C1 vectors using the cut sites BglII/SalI (vector) and BamHI/XhoI (insert). SetA-CTD mutants used in this study were generated using site-directed mutagenesis protocols adapted from the Qiagen QuikChange II Site-Directed Mutagenesis Kit. All constructs were verified by DNA sequencing.

### Protein Expression and Purification

All SetA 523-629 constructs (untagged SetA mutants and GFP-SetA fusion constructs) in pET28(a) His-Sumo were transformed into the Rosetta (DE3) strain of *E.coli* cells using the antibiotic selection markers kanamycin and chloramphenicol. Transformed bacterial cells were grown in 1L expression cultures at 37°C at 220 rpm and induced with 0.25 mM IPTG during log-phase growth (A_600nm_ 0.6-0.8). Cells were incubated at 18°C and 180 rpm for 18 hours post-induction. Cells were collected by pelleting expression cultures at 4500 rpm for 15 minutes at 4°C. Cells were resuspended in 20 mL of 50 mM tris, 150 mM NaCl pH 7.5 containing 1 mM PMSF. Cells were lysed by two rounds of sonication at 50% amplitude, 2-minute duration, 2 sec on/off pulse on ice. Sonicated samples were spun at 16,000 rpm for 45 minutes at 4°C to remove the insoluble fraction. The supernatant was collected and mixed with 2 mL of cobalt resin and incubated while rotating for 4 hours at 4°C to bind proteins. Protein-bound resin was washed with several column volumes of 50 mM tris, 150 mM NaCl pH 7.5 to remove unbound and nonspecifically bound proteins. The resin was resuspended in 3 mL of wash buffer and cut overnight with His-tagged Ulp1 at 4°C. Cut proteins were eluted the next day and concentrated to a final volume of 3 mL using a 30 kDa cut-off centrifugal concentrator. Proteins were run on a Superdex200 16/200 column using an AKTA GE Healthcare FPLC system. Peak fractions were run on SDS-PAGE, and SetA protein fractions were collected and pooled. Purified proteins were further concentrated and stored at −80°C. For NMR protein sample preparation, the isotopically labeled protein was produced using the appropriate Spectra-9 bacterial growth medium (SpetraStableIsotopes). The proteins were purified similarly as described above. The final samples were buffer exchanged to 20 mM sodium phosphate, pH 6.0, with 0.05% azide, 5% ^2^H2O, 10 μM 2,2-dimethyl-2-silapentane-5-sulfonate, and 1 μM of the protease inhibitors PMSF.

### Protein NMR Spectroscopy

All NMR spectra were collected at 25 °C on a Varian INOVA 500 MHz spectrometer and used pulse sequences available in the Varian BioPack user library. Sequential and aliphatic side chain assignments were determined by analysis of 3D-HNCO, HN(CO)CA, HNCA, HNCACB, HCACO, CBCA(CO)NH, and HCCH-TOCSY NMR experiments. All spectra were processed with NMRPipe (56) and subjected to visual analysis in Sparky (57). NOE correlations between nearby protons were identified in 3D ^15^N-NOESYHSQC, ^13^C-NOESYHSQC (aromatic), and ^13^C-NOESYHSQC (aliphatic) NMR spectra. NMR structure calculation was performed via Ponderosa-C/S program package (46). Briefly, raw NOESY data, chemical shift assignments, and the protein sequence were submitted through Ponderosa Client and the initial structure was calculated by the Ponderosa Server. The calculated structure ensembles and constraints were analyzed via Ponderosa Analyzer. Hydrogen bonds between β-strands and within α helices were identified during a later stage of structural refinement and were added to the later cycles of structural calculation. During the iterative refinement cycles, violated NOE restraints were loosened, deleted, or reassigned based on the experimental NOE spectra. The final 20 structures with the lowest target function values from 200 independently calculated structures were retained as the final ensemble. Structure representations were drawn using PyMol (DeLano Scientific LLC).

### Cell culture, transfection, and fluorescent microscopy

For co-localization analysis, mCherry-SetA-CTD constructs were transiently co-expressed with an EGFP-2xFYVE domain construct in HeLa cells. Cells were passaged at 25-30% initial density in a 24-well plate in D10 media. Cells were subsequently transfected 24-hours later with 0.15 μg of each plasmid and a 1:1 (m/v) ratio of Lipofectamine 2000 in Opti-MEM for a total volume of 50 μL. At 14-16 hours post-transfection, cells were fixed in 4% paraformaldehyde in PBS solution for 20 minutes at room temperature then washed three times in PBS. Fixed coverslips were mounted onto glass slides using Fluoromount-G mounting solution. Fixed cells were imaged using a spinning disk confocal microscope (Intelligent Imaging 108 Innovations, Denver, CO) equipped with a spinning disk confocal unit (Yokogawa CSU-X1), an inverted 109 microscope (Leica DMI6000B), a fiber-optic laser light source, a 100× 1.47NA objective lens, 110 and a Hamamatsu ORCA Flash 4.0 v2+ sCMOS camera. Images were acquired and processed using the Slidebook (version 6) software.

The colocalization of SetA-CTD with the 2xFYVE domain was quantified using ImageJ software. To determine the percent SetA co-localized with 2xFYVE, the total SetA CTD signal was measured and defined as “total SetA”. The fraction of SetA CTD co-localized with the 2xFYVE domain was measured by quantifying the mCherry-SetA signal using a “mask” generated from the 2xFYVE domain (GFP) channel. Percent colocalization was calculated by dividing the co-localized mCherry-SetA-CTD signal by the total mCherry-SetA-CTD signal. Measured signal intensities were normalized to the background for each image analyzed.

### Giant Unilamellar Vesicles (GUVs) binding assay

POPC (1-palmitoyl-2-oleoyl-*sn*-glycero-3-phosphocholine) and POPS (1-palmitoyl-2-oleoyl-sn-glycero-3-phospho-L-serine) were purchased from Avanti Polar Lipids (Alabaster, AL, USA). DiD (1,1’-Dioctadecyl-3,3,3’,3’-Tetramethylindodicarbocyanine, 4-Chlorobenzenesulfonate salt) was purchased from ThermoFisher scientific (Molecular probes) (Waltham, MA, USA). Dye concentrations were measured in methanol by absorbance spectroscopy. We used molar extinction coefficient 91800 (M.cm)^-1^.

Giant Unilamellar Vesicles (GUVs) were prepared using the electroformation procedure, with a few modifications. Lipids and dyes were diluted in chloroform and were gently spread on glass slides covered with ITO (Indium-Tin-Oxide) to form a lipid film on the slide. The lipid film was hydrated with a 100 mM sucrose solution at 55 °C. an oscillating potential of 1.4 V and 12 Hz was subsequently applied for 3 hours in the ITO slides to form the GUVs. The temperature controller was set to slowly cool the GUVs to room temperature at a rate of 2 °C/hour and then diluted the GUVs in a buffer of 95 mM Tris (2-Amino-2-(hydroxymethyl) propane-1,3-diol), pH = 7.4. We adjusted the osmolarity of the buffer and the sucrose solution to 100 mOsm. GUVs were prepared with the lipid compositions POPC/POPS/PI(3)P = 0.85/0.10/0.05 or POPC/POPS = 0.90/0.10, and labeled with the fluorescence dye DiD with a ratio of dye/lipid = 1/2500 (0.04 mol%).

To test the protein binding in these GUVs, 2 μL of GUVs and 2 μL of each EGFP-tagged protein (1 mg/ml) were added to a microscope slide and incubated for ?? minutes. these samples were then imaged using a confocal microscope Nikon Eclipse C2+ (Nikon Instruments, Melville, NY), with objective (60X, N.A. 1.2). The fluorescence signal intensity of EGFP on the GUV counter was measured using a script built-in ImageJ (Fiji). For each experimental condition, the final value is averaged from 10 GUV measurements.

## Acknowledgments

We thank Dr. Gerald W. Feigenson (Cornell University) for technical help in GUV binding assay. This work was supported by National Institutes of Health (NIH) Grants RO1-GM135379-01 (Y.M.).

## Author contributions

Y.M. conceived the project. W.H.J.B., T.A.E., X.W., Q.Z., R.E.O., and Y.M. performed the experiments. W.H.J.B., T.A.E., L.K.N., R.E.O., and Y.M. analyzed the data. W.H.J.B. and Y.M. wrote the paper.

---

**Supplemental Figure 1. Multiple sequence alignment of the SetA-CTD domain.** The sequences corresponding to the SetA-CTD domain (aa. 522-629) from different *Legionella* species were aligned by Clustal Omega and colored using Multiple Align Show (http://www.bioinformatics.org/sms/index.html). SetA residue numbers are marked above the alignment with secondary structural elements drawn above. Identical residues are highlighted in dark grey and similar residues in light grey. Secondary elements are drawn above the alignment. Loop 1 and Lop 2 loop are marked with squares. R612 is marked by “&”. Membrane insertion hydrophobic residues (L549, L550, F608, and F609) are highlighted with “*”. NCBI accession numbers are as follows: SetA, *Legionella pneumophila*, AAU28047.1; lgra0302, *Legionella gratiana*, WP_058498745.1; lpp1961, *Legionella pneumophila Paris*, CAH13113; lpl1955, *Legionella pneumophila Lens*, CAH16195; Lstei2448, *Legionella steigerwaltii*, WP_058477388.1; LmasA1472, *Legionella massiliensis*, WP_043873545.1; Lwad2150, *Legionella wadsworthii*, WP_031567619.1; Lpar0355, *Legionella parisiensis*, WP_058518790.1; LboB3024, *Legionella bozemanae*, WP_131780238.1; Ltuc2674, *Legionella tucsonensis*, WP_058520732.1; Lche0549, *Legionella cherrii*, WP_028380196.1.

**Supplemental figure 2. ^1^H-^15^N HSQC spectrum of SetA-CTD.** The HSQC spectrum of ~1 mM ^15^N SetA-CTD in 20 mM sodium phosphate (pH 6.0) recorded at 25 °C and 500 MHz. Peaks are labeled with the one-letter amino acid code and sequence number.

**Supplemental Figure 3. Distribution of NOE distance constraints distribution for each residue in SetA-CTD used in structure calculation.** Low range (i-j <=1) distance constraints are colored in white, medium range (1< i-j <=4) constraints are shown in salmon, and long-range (4 < i-j) constraints are in blue.

