## Supplementary figures and images for "Solution structure of the phosphatidylinositol 3-phosphate binding domain from the *Legionella* effector SetA"

### supplemental figure 1

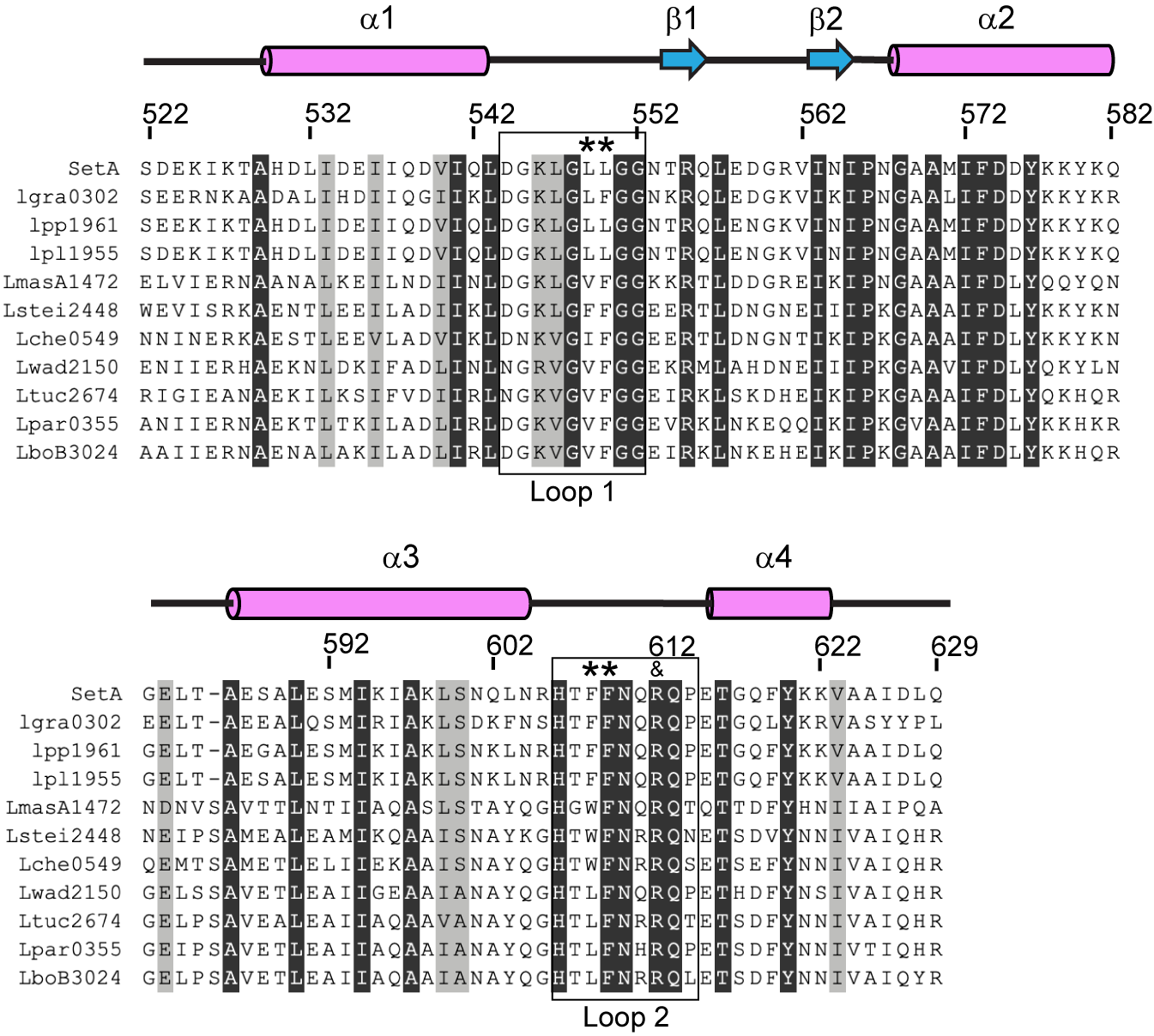

### supplemental figure 2

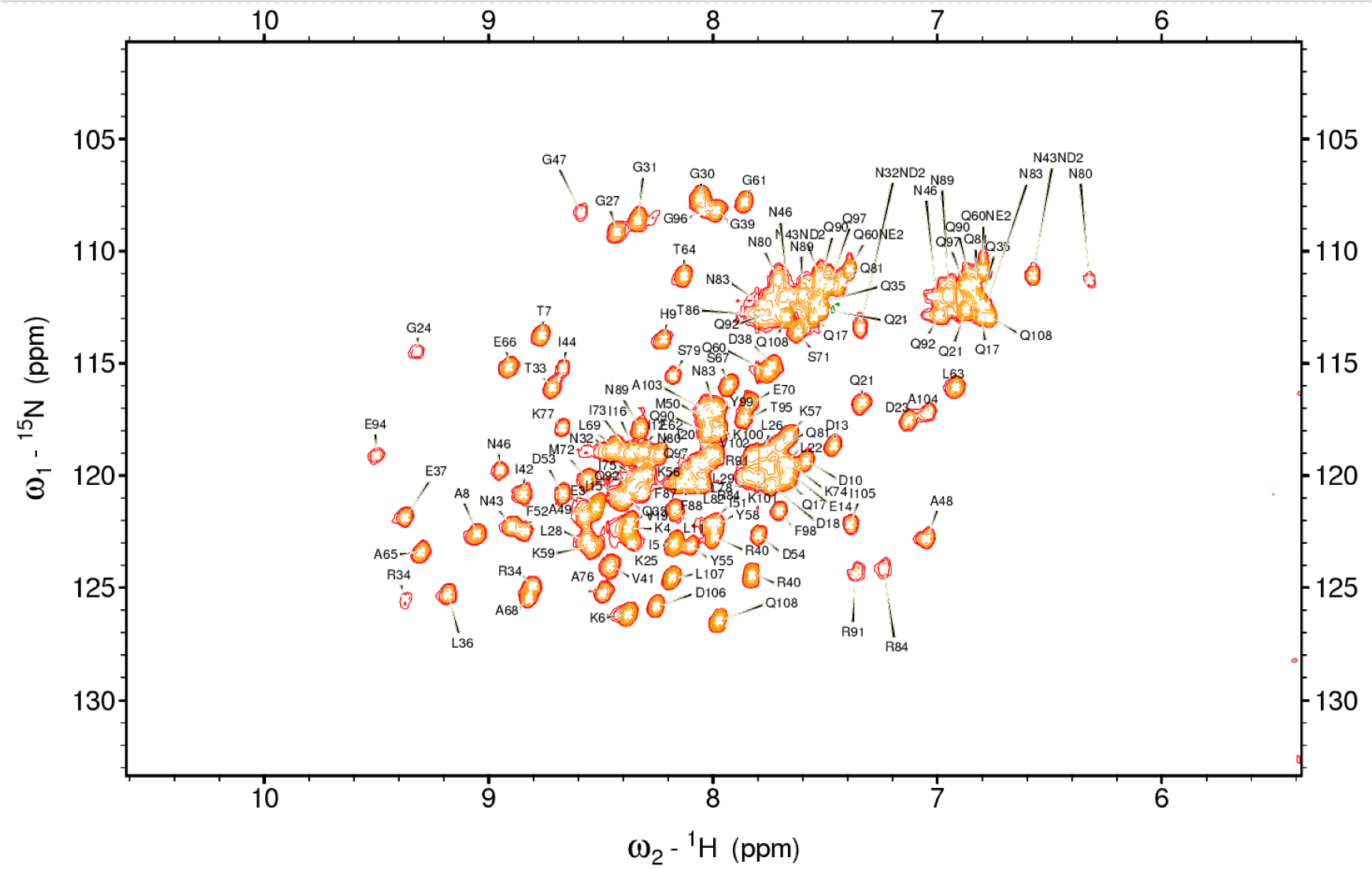

### supplemental figure 3

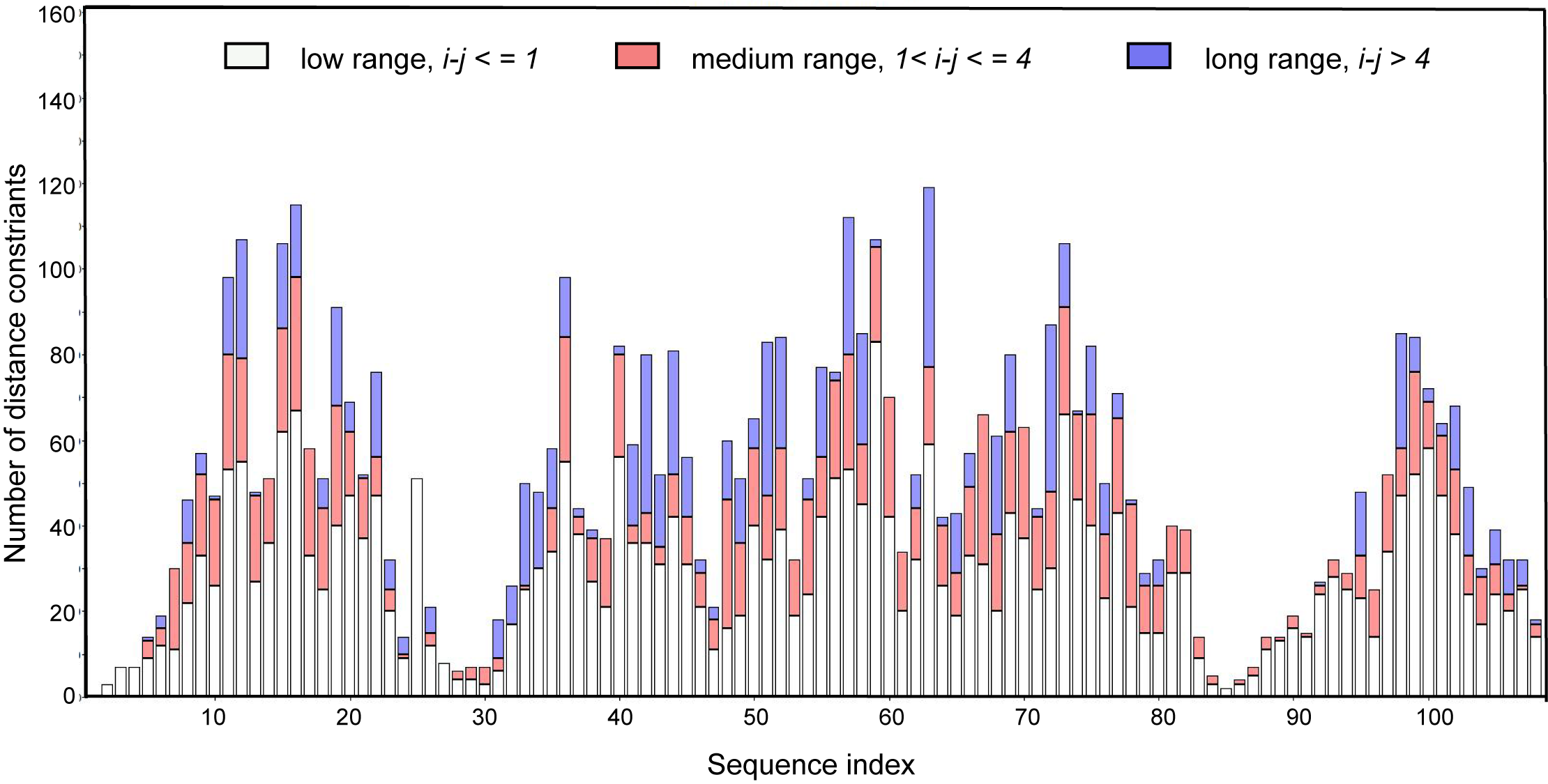
